# Cell-Level Virtual Screening

**DOI:** 10.64898/2026.05.11.724149

**Authors:** Caleb N. Ellington, Sohan Addagudi, Jiaqi Wang, Benjamin Lengerich, Eric P. Xing

**Author notes:** Equal contribution (Caleb N. Ellington, Sohan Addagudi, Jiaqi Wang).

## Abstract

Virtual screening methods prioritize therapeutic candidates by predicting molecular properties and interactions. However, molecular models are insufficient to predict higher-order effects that arise in real biological systems, leading to late-stage failures in drug discovery. Virtual cells have been posed as a solution to this problem by predicting gene expression responses to drugs, but they remain weakly validated as screening tools; gene expression is only an intermediate in understanding drug success or failure. Despite burgeoning progress in virtual cells, some basic questions remain. Is expression even a good representation of higher-order drug effects? How can expression and other cell-level representations be applied to prioritize therapeutic candidates? Can cell-level methods be fairly compared against traditional molecular-level screens? We address these questions in a two-pronged approach. First, we curate two benchmarks, Drug-Disease Retrieval Bench (DDR-Bench) and Drug-Target Retrieval Bench (DTR-Bench), which directly compare cell-level methods against traditional molecular methods on canonical drug discovery tasks. DDR-Bench evaluates a method’s ability to prioritize disease indications for drugs with novel target profiles. DTR-Bench evaluates a method’s ability to reconstruct drug-target interactions from separate perturbation modalities that act on shared mechanisms, bridging the gap between cell-level methods and classic molecular screens. We identify shortcomings of existing screening methods on these benchmarks, and propose an alternative representation of drug effects: perturbed gene networks. Inferring post-perturbation gene networks on-demand for unseen drugs requires methods that generalize beyond traditional plug-in network estimators. We develop a scalable differentiable surrogate loss for multivariate Gaussians, which we apply to train a context-adaptive amortized estimator that maps perturbation metadata to gene-gene dependency network parameters. The resulting model, CellVS-Net, achieves SOTA on predicting how gene networks restructure under a variety of complex multivariate experimental conditions, including different cell types, small molecule therapeutics, signaling molecules, gene knockdowns, and gene over-expressions. When compared to other molecular and cell-level representations of drugs, we find that CellVS-Net achieves SOTA on both virtual screening benchmarks. Overall, CellVS-Net demonstrates that cell-level virtual screening methods are a viable alternative to molecular screening, and associated benchmarks enable hill-climbing on relevant drug discovery tasks.

## Introduction

Despite continuous improvements in virtual screening for molecular interactions, recently achieving near-instantaneous proteome-scale screens [1], virtual screening approaches still remain blind to emergent failures at the level of cellular systems. Moving beyond molecular interactions, large-scale expression and morphology profiling efforts such as LINCS L1000 [2], Tahoe-100M [3], scPerturb [4], JUMP-CP [5], and Recursion’s phenomics platform [6] aim to support phenotype-level screens by experimentally measuring cell states under many perturbations and ranking candidates by similarity or reversal of disease signatures. However, these phenotypic screens require running a new assay for each candidate drug and are therefore limited by experimental throughput and design. In contrast, virtual screening aims to predict cell-level responses for unseen perturbations based only on their molecular or target features, enabling *in silico* ranking of large candidate libraries before any experiment is performed. To be practically useful, such a framework must both capture system-level cellular responses and generalize reliably to drugs, targets, and contexts that were never observed.

A growing number of works on virtual cells aim to extend these efforts by training machine learning models to simulate cellular behavior in response to perturbations [7, 8]. To the best of our knowledge, all methods that predict cellular response to unseen perturbations are primarily evaluated on their ability to reconstruct a post-perturbation readout (e.g. expression, morphology, IC50) [9, 10, 11, 12, 13, 14, 15, 16, 17, 18, 19, 20]. However, expression (predicted and real) is an intermediate representation of perturbation effect, and does not directly identify the safety and efficacy of a therapeutic. Miladinovic et al. [21] go beyond reconstruction-based evaluations to investigate synonymous genetic and pharmacological perturbations, but this method is unable to generalize to perturbations beyond its training vocabulary. To enable cell-level virtual screening, we believe that the next generation of virtual cell methods must (i) provide insights directly relevant to drug discovery and (ii) generalize arbitrarily to unseen therapeutics to promote virtual exploration. To address this, we curate the Drug-Disease Retrieval Bench (DDR-Bench) and Drug-Target Retrieval Bench (DTR-Bench) benchmarks. DDR-Bench evaluates whether a method can identify the correct FDA-approved disease indication for a drug with a novel target profile; a failure case for target-based screening that we hypothesize requires a system-level model of drug response. DTR-Bench extends a traditional drug discovery task (drug-target interaction prediction) to cell-level screening. While molecular methods often predict these interactions directly, cell-level methods should identify synonymous small molecule and gene knockout perturbations based on the similarity of cell-level responses. We assemble a representative panel of molecular and cell-level representation methods for screening, and identify shortcomings on both DDR-Bench and DTR-Bench.

Based on these shortcomings, we also investigate alternatives for cell-level drug effect representation. In particular, we hypothesize that post-perturbation gene networks would capture how perturbations rewire cellular circuitry, providing a richer description of perturbation effects than expression snapshots. However, most network inference methods rely on plug-in estimators with large cohorts [22, 23], which fail to capture the continuous and context-dependent rewiring and cannot generalize to new perturbation conditions, a key requirement for virtual screening.

To enable in-context prediction of network rewiring, we develop a scalable differentiable surrogate for the multivariate Gaussian negative log-likelihood that decomposes covariance estimation into pairwise regression problems. This avoids direct optimization of the expensive log |Σ| and Σ^−1^ terms in Gaussian likelihood, and serves as a drop-in replacement for the isotropic-Gaussian (mean squared error) losses commonly used in cellular response models. We apply this new objective to train CellVS-Net, a model which maps multivariate cellular and therapeutic contexts (cell type, drug, dose) to predict a post-perturbation gene network represented as a Gaussian graphical model. Rather than fitting a separate network for each cell-drug–dose combination, CellVS-Net learns a single context encoder that functions as an amortized estimator for gene networks, going directly from data context to network parameters. Critically, this allows us to generate networks in-context without requiring context-specific data collection. We train CellVS-Net to generate networks for unseen cell lines and perturbations including small molecule therapeutics, signaling molecules, gene knockdowns, and gene over-expressions, and apply the network predictions to the zero-shot DDR and DTR benchmarks.

Our contributions are as follows:

1. We provide the first evaluation of virtual cell methods’ ability to prioritize drugs in practical virtual screening settings, assembling a representative panel of molecular and cell-level methods for fair cross-modal comparison. We develop DDR-Bench, which evaluates whether a method can identify correct disease indications for drugs with novel target profiles, and DTR-Bench, which extends drug-target interaction modeling to cell-level methods, bridging the conceptual gap between molecule-level and cell-level screens.
2. We identify shortcomings of virtual cell methods on these benchmarks, motivating new representations of cell-level drug effects.
3. We posit perturbed gene networks as an improved representation of cell state and develop a scalable differentiable surrogate loss for multivariate Gaussians to train CellVS-Net, a context-adaptive amortized estimator for generating gene networks on-demand.
4. CellVS-Net accurately maps multivariate contexts including cell types, small molecule therapeutics, signaling molecules, gene knockdowns, and gene over-expressions to context-specific gene dependency networks.
5. CellVS-Net achieves SOTA on predicting networks for held-out perturbations. When applied as a screening representation, CellVS-Net also achieves SOTA on the zero-shot DDR and DTR benchmarks.

## Methods

### Benchmarks

To evaluate and develop cell-level virtual screening methods, we first curate a comprehensive database containing drug, target, expression, and disease approval data by combining the OpenTargets and LINCS databases. OpenTargets provides structured databases of drug-disease pairs based on Phase-IV FDA approval data, drug-target pairs based on known mechanisms of action, and target-disease associations from various lines of evidence. LINCS provides post-perturbation gene expression data for a variety of perturbation types including small molecule therapeutics, signaling molecules, gene knockdowns, and gene over-expressions for multiple cell lines. By merging these sources, we produce a large database of (perturbation, target, disease, expression) for multiple perturbation types, which can be reused for evaluation of molecule-based, target-based, and cell-based screening methods.

### Representing Drug Effects

Both benchmarks reduce each perturbation to a fixed-length vector and compare drugs or targets by Euclidean distance. This is deliberately modality-agnostic: any vector representation (e.g. expert-derived molecular features, foundation-model embeddings, or our predicted networks) plugs into the same retrieval and edge-classification protocols; no sample splitting is required; and the protocol scales as new drugs receive approval, mirroring billion-scale semantic search [1]. This provides a stringent zero-shot evaluation setting in which performance differences are interpreted entirely in terms of how well each representation captures therapeutically meaningful cell-level effects.

### Drug-Disease Retrieval

DDR-Bench evaluates whether a method can identify the correct FDA-approved disease indication for a drug with an unseen target profile; a failure case for target-based screening that we hypothesize requires a system-level model of drug response. Let 𝒟 denote the set of curated drugs and let each drug *q* ∈ 𝒟 be represented by a fixed-length vector *ϕ*_*q*_ ∈ ℝ^*d*^ produced by the method under evaluation (e.g., a CellVS-Net network, a fingerprint, or an expression embedding); the embedding dimension *d* is method-specific. For a query drug *q* ∈ 𝒟, the reference set ℛ = 𝒟 \ {*q*} consists of all other drugs with different target profiles. We rank the references in ascending order of Euclidean distance 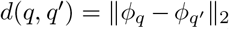 for *q*^*′*^ ∈ ℛ. Let *y*^⋆^(*q*) denote the FDA-approved disease indication of *q*, drawn from a finite set of indications 𝒴, and let *y*_(1)_, …, *y*_(|ℛ|)_ ∈ 𝒴 denote the indications of the ranked reference drugs (so *y*_(*r*)_ is the indication of the *r*-th closest reference drug). We define

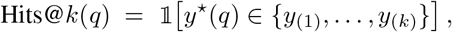

where 𝟙 [·] is the indicator function returning 1 when the condition holds and 0 otherwise, and report the mean over queries for *k* ∈ {1, 5, 10, 25} (Figure 2). Dataset construction details are deferred to Appendix E.

**Figure 1:**
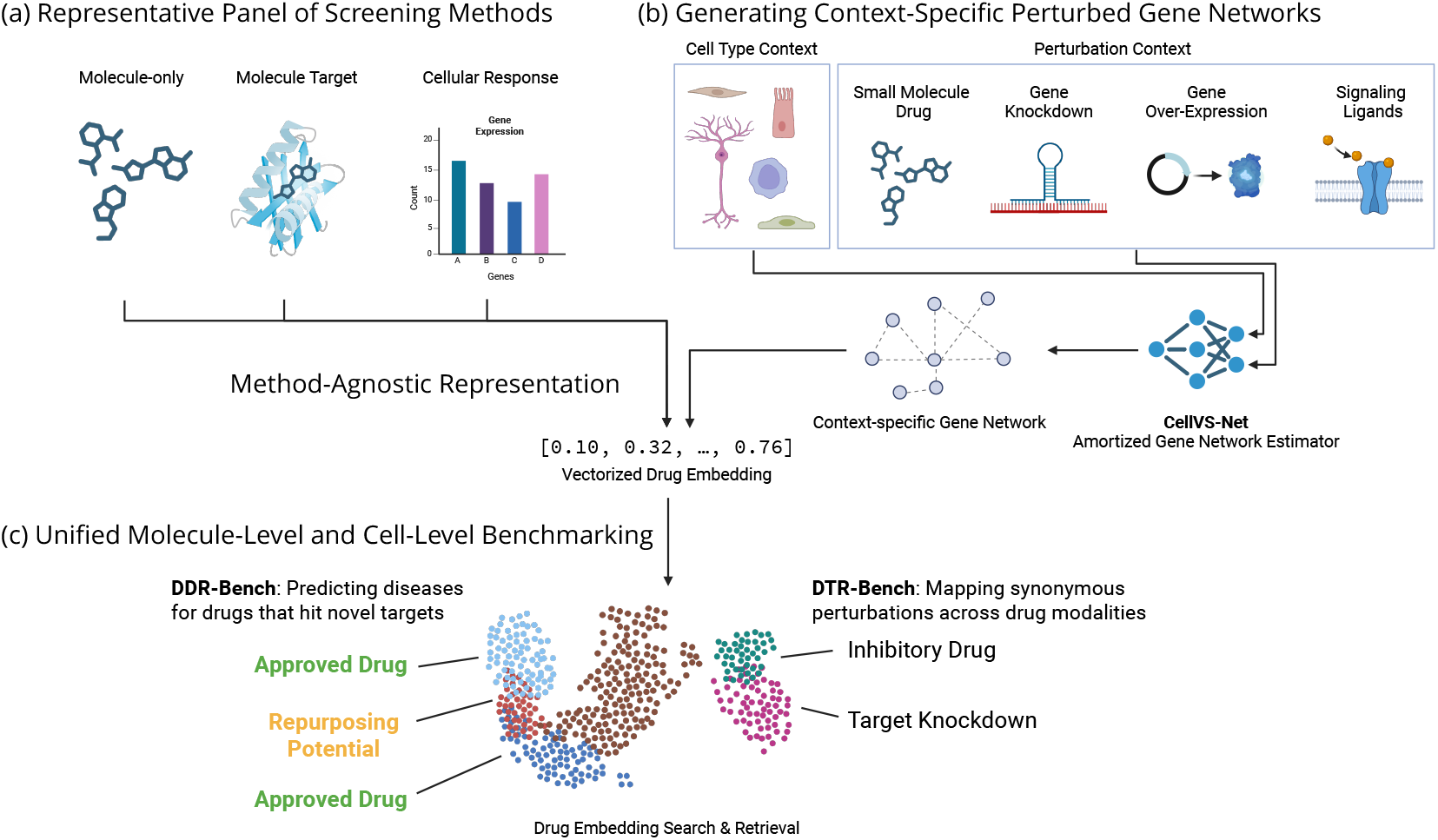
Graphical abstract. (a) We assemble a representative panel of screening methods for representing drug effects at molecule, target, or cell level. (b) We propose gene networks as an improved representation of cell-level drug effects, and develop CellVS-Net, a model that generates context-specific perturbed gene networks on-demand for unseen drugs and cell types. (c) To fairly compare molecule, target, and cell-level representations of drug effects, we introduce two modality-agnostic benchmarks for evaluating screening methods: DDR-Bench and DTR-Bench.

**Figure 2:**
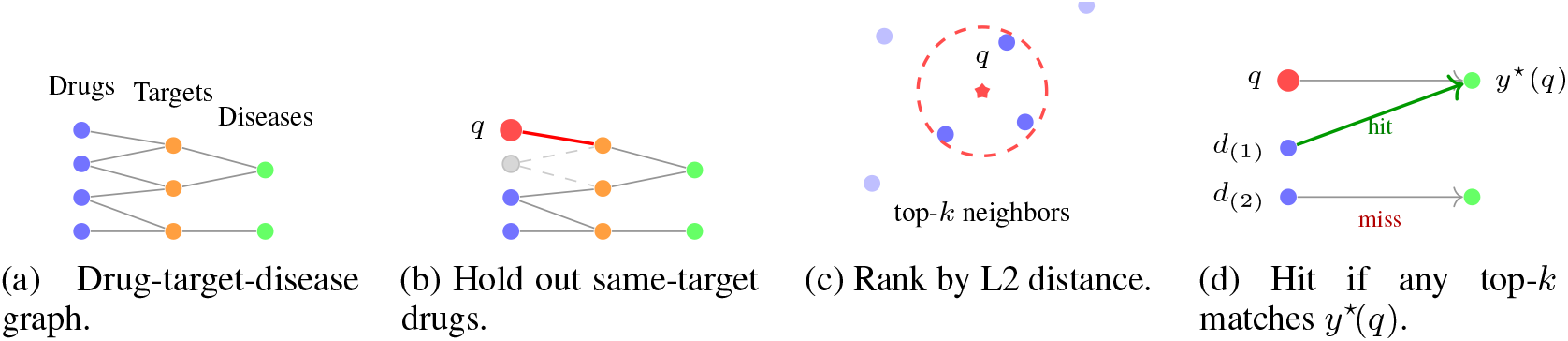
DDR-Bench evaluation. (a) Drugs, targets, and diseases form a tripartite graph. (b) For each query drug *q*, all reference drugs sharing *q*’s target signature are masked. (c) The remaining reference set is ranked by L2 distance to *q* in the chosen representation space. (d) Each retrieved drug *d*_(*i*)_ is paired with its FDA-approved disease; a top-*k* neighbor is a hit when its disease equals the query’s FDA-approved disease *y*^⋆^(*q*), and a miss otherwise. Hits@*k* is 1 when at least one of the top-*k* neighbors is a hit.

### Drug-Target Retrieval

DTR-Bench reconstructs known drug-target interactions from cell-level perturbation signatures, bridging molecular and cell-level screens. Let 𝒟 denote the set of small-molecule drugs (LINCS chemical perturbations) and 𝒯 the set of protein targets (LINCS shRNA knockdowns of those targets); each drug *i* ∈ 𝒟 and each target *j* ∈ 𝒯 is mapped to a vector *ϕ*_*i*_, *ϕ*_*j*_ ∈ ℝ^*d*^ in a shared representation space (the embedding dimension *d* is method-specific). Let *d*_*ij*_ = ∥*ϕ*_*i*_ − *ϕ*_*j*_∥_2_ denote their Euclidean distance and *y*_*ij*_ ∈ {0, 1} the ground-truth interaction label, with *y*_*ij*_ = 1 iff drug *i* binds target *j* in the curated drug-target graph. We sweep a threshold *τ* ∈ ℝ_+_ on *d*_*ij*_ over all |𝒟| × |𝒯| pairs, predicting *ŷ*_*ij*_(*τ*) = 𝟙 [*d*_*ij*_ ≤ *τ*], and report AUROC and AUPRC for {*ŷ*_*ij*_(*τ*)} versus {*y*_*ij*_} as *τ* varies. We additionally perform bidirectional Hits@*k* retrieval: for a drug query *i* ∈ 𝒟, we rank all targets *j* ∈ 𝒯 in ascending order of *d*_*ij*_ and define

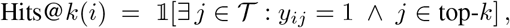

with the symmetric definition for target queries (ranking drugs *i* ∈ 𝒟 for a target query *j* ∈ 𝒯). We report Hits@*k* for *k* ∈ {1, 5, 10, 50}. Figure 3 illustrates the protocol. Different drug modalities are measured with different assays, so expression-based representations are susceptible to batch effects; before evaluation we PCA-decompose the combined drug and target representations and remove up to the first 3 principal components, reporting the best of the 4 attempts (removing 0, 1, 2, or 3 components). Dataset construction details are deferred to Appendix E.

**Figure 3:**
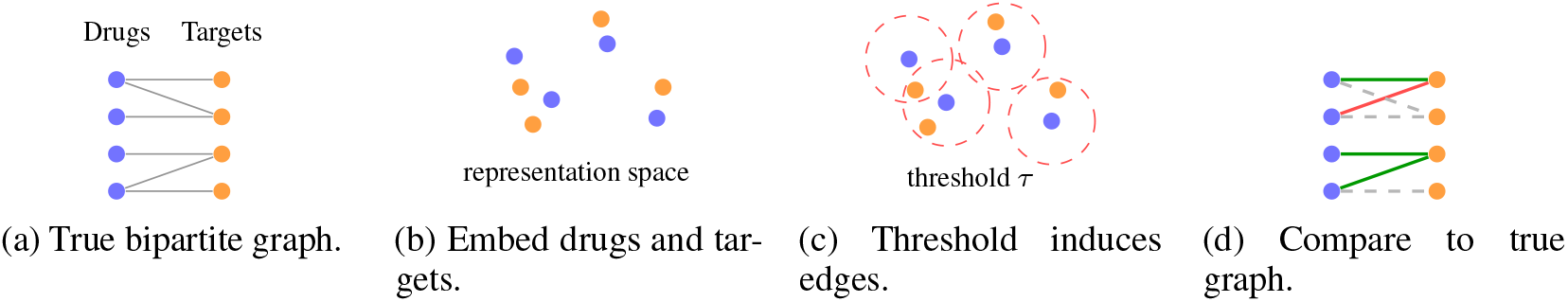
DTR-Bench evaluation. (a) Ground-truth drug-target interactions form a bipartite graph. (b) Each method maps drugs and targets into a shared representation space. (c) A distance threshold *τ* on *d*_*ij*_ = ∥*ϕ*_*i*_ − *ϕ*_*j*_∥_2_ induces a predicted bipartite graph; AUROC/AUPRC are obtained by sweeping *τ*. (d) Comparing predicted to true edges yields true positives (solid green), false positives (solid red), and missed true edges (dashed grey); Hits@*k* is computed by ranking targets per drug and vice versa.

### Representative Panel of Screening Methods

#### Drug-only

For a cell and target-agnostic baseline, we include a molecular fingerprint baseline [24]. Circular fingerprints remain a workhorse for classical virtual screening pipelines and ligand-based similarity search. Including this baseline grounds our evaluation against a mature, purely structure-based representation that does not see any cellular readout or target information.

#### Drug-target Interactions

We include SPRINT [1], a recent proteome-scale ligand-protein binding model with strong performance on drug-target interaction prediction. For each drug, we use the SPRINT-predicted binding profile against a fixed reference proteome as its vector representation. This baseline is directly optimized for drug-target interaction prediction and grounds our evaluation against a strong target-aware molecular method that uses no cellular readout.

#### Gene Expression

Most virtual cell methods predict post-perturbation gene expression from compound structure. However, existing methods are not capable of generalizing to the breadth of held-out perturbations considered here. To represent expression reconstruction methods, we construct an expression-prediction baseline that takes a ChemBERTa embedding of a drug’s SMILES and regresses to either the LINCS L1000 landmark expression vector (Appendix D). The predicted expression then serves as the drug’s representation.

#### Cell Embedding

We also include two expression embedding baselines to capture latent cell states. Embedding-based methods can outperform other methods on these tasks by removing redundant expression features and distilling cell states into semantically meaningful low-dimensional latent features. As a simple strong baseline, we compare PCA-compressed expression, which provides semantically meaningful low-dimensional features through simple linear compression. We also assess foundation model (FM) embeddings using AIDO.Cell [11]. To predict these embeddings for held-out therapeutics where we don’t have observed expression data, we adopt a latent prediction approach where we train a multi-layer perceptron to predict the latent features of the post-perturbation expression embedding.

#### Oracles

In a realistic virtual screening scenario, expression measurements will *not* be available: the goal is precisely to avoid running large numbers of physical experiments. In this analysis, we also include observed post-perturbation expression as an “oracle”, assessing the value of real data acquisition. We can conceptually treat observed expression as the output of an idealized virtual cell that is a perfect generator of the true transcriptional response. Any method that predicts expression is ultimately trying to approximate this oracle. Using expression as a baseline therefore serves two purposes: (i) it provides an optimistic upper bound for expression-based objectives. If a task is hard even with the true expression snapshot, no virtual cell that only regresses expression can be expected to perform substantially better; and (ii) it lets us ask whether other representations based on idealized expression can *surpass* the utility of raw expression snapshots. We also consider “oracle” embeddings (both PCA and FM) by directly applying these methods to the oracle expression.

### Cell-level Screening with Gene Networks

We identify shortcomings with both molecular and cell-level methods on DDR-Bench and DTR-Bench. To go beyond expression snapshot prediction and embedding, we propose representing drug effects using post-perturbation gene networks. This requires the development of a scalable method for amortized estimation, which can produce gene networks on-demand for unseen perturbations.

#### Notation and setup

Each biological sample *n* is a *p*-dimensional gene-expression vector *X*_*n*_ ∈ ℝ^*p*^ (*X*_*n,i*_ is the expression of gene *i*), paired with a context vector *C*_*n*_ ∈ ℝ^*m*^ that encodes its experimental conditions (cell-type identity, perturbation modality, perturbation payload content, dose, and measurement time). We model samples as draws from sample-specific distributions *X*_*n*_ ∼ *P* (*X* | *θ*_*n*_), where *θ*_*n*_ contains all distribution parameters (mean, variance, dependence structure).

To share information across samples, we treat parameters as a function of context,

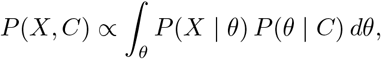

where the context encoder *P* (*θ* | *C*) is our desired amortized estimator. Following the contextualized modeling framework [25], we implement the context encoder as a deterministic deep network *f* : ℝ^*m*^ → Θ giving *P* (*θ* | *C*) = *δ*(*θ* − *f* (*C*)) (with *δ* the Dirac delta) and *P* (*X*_*n*_ | *θ*_*n*_) = *P* (*X*_*n*_ | *f* (*C*_*n*_)). This regime has been extended to linear models [26, 27, 28] and several graphical model classes [29, 30], but the general multivariate Gaussian case remains unaddressed.

#### Multivariate Gaussian loss

We seek a contextualized multivariate Gaussian, *X* |*C* ∼ 𝒩 (*µ*(*C*), Σ(*C*)), with mean vector *µ*(*C*) ∈ ℝ^*p*^ and covariance matrix Σ(*C*) ∈ ℝ^*p*×*p*^, Σ(*C*) ≻ 0, both produced by neural encoders, and optimized end-to-end with stochastic gradient descent. Up to constants, the exact negative log-likelihood for one sample is

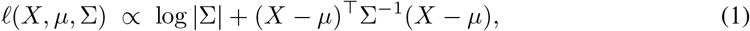

where log |Σ| denotes the log-determinant of Σ. Optimizing this objective requires recomputing Σ^−1^, log |Σ|, and a positive-definite constraint at every gradient step, which is prohibitive to run on every sample batch even for small gene panels (*p* ∼ 50). We instead optimize a composite-likelihood surrogate built from the regression form of Pearson’s correlation between genes *i* and *j*,

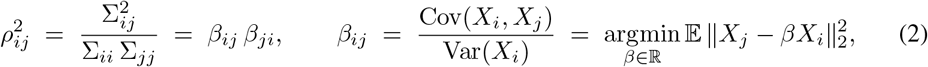

where Σ_*ij*_ is the (*i, j*) entry of Σ, *β*_*ij*_ ∈ ℝ is the ordinary least-squares (OLS) coefficient for regressing gene *X*_*j*_ on gene *X*_*i*_, and *β* ∈ ℝ^*p*×*p*^ collects all pairwise coefficients. Let 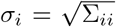 denote the marginal standard deviation of gene *i*, with 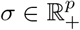 the vector of all marginal standard deviations and *µ* ∈ ℝ^*p*^ the marginal mean vector. The marginalization properties of Gaussians let each (*i, j*) pair be fit independently and reassembled via

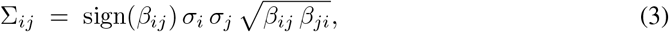

where *µ* and *σ* are estimated under the per-gene isotropic Gaussian likelihood 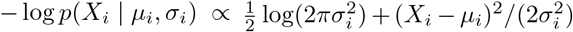. The full contextualized objective for one sample with context *C* is

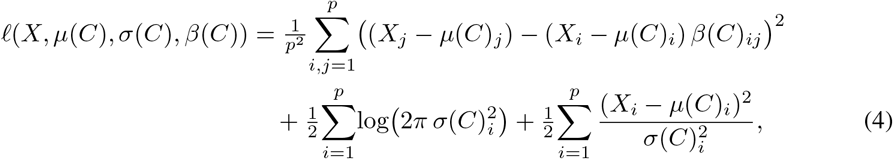

which is differentiable, jointly fits the context encoders *µ*(*C*), *σ*(*C*), *β*(*C*), and avoids any explicit matrix inversion or determinant. The estimated parameters yield a context-specific Gaussian graphical model when the induced correlation matrix is positive definite, and project onto the positive-definite cone otherwise. This surrogate is a pseudo-likelihood rather than the exact NLL, but inherits the interpretability of Gaussian covariance and serves as a drop-in replacement for the mean-squared-error losses used in current cell models; we report the pairwise regression term alone as MSE.

#### CellVS-Net

Drug efficacy, toxicity, and mechanism of action emerge from coordinated shifts in gene–gene dependencies rather than from marginal expression alone. We therefore seek a representation that describes the system-level effect of a perturbation and generalizes to unseen drugs, targets, and cell types. We apply this new loss to create CellVS-Net, a deep learning model that learns to generate Gaussian graphical models representing gene networks on-demand for unseen drugs, targets, and cell types, capturing context-specific network restructuring. With CellVS-Net, each perturbation is represented not by its transcriptomic snapshot but by the inferred structure of gene–gene relationships that best explains the observed data under that context. The context encoder accommodates many different biochemical representations of a perturbation; modality-specific encoders for small molecules, protein targets, and genetic perturbations are detailed in Appendix C. Implementation details on the architecture and optimizers used are discussed in the Appendix.

We use CellVS-Net’s predicted gene network as a structured cell-level representation. Each drug is represented by the upper-triangular squared correlations *β*_*ij*_ (*C*)*β*_*ji*_ (*C*) concatenated with the predicted mean shift *µ*(*C*), capturing coordinated rewiring of gene-gene dependencies rather than marginal expression alone.

## Results

### Generating Perturbation-specific Gene Networks

In order to apply post-perturbation gene networks to screening, we first consider the problem of estimating perturbation-specific gene networks. Traditional plug-in estimators fit an independent network for each cell line or perturbation. This approach overfits severely in low-sample regimes and cannot produce estimates for unseen conditions. Population models avoid overfitting, but collapse all samples from these heterogeneous contexts into a single model. To enable virtual screening, we require a network estimator which is adaptable to completely unseen contexts and perturbations. CellVS-Net addresses this by learning a smooth mapping from context features to network parameters. We train two variants: CellVS-Net Target and CellVS-Net Molecule. CellVS-Net Target takes as input a target gene or protein, a basal unperturbed expression profile for the cell line of interest, and a perturbation modality indicator (one of chemical, knockdown, over-expression, of ligand). CellVS-Net Molecule only operates on chemical perturbations, and takes as input a small molecule SMILES and a basal unperturbed expression profile for the cell line of interest.

In experiments across perturbation modalities, CellVS-Net reduces network MSE by 47% on chemical perturbations and by 12–37% on the genetic and ligand modalities relative to the population baseline (Table 1). CellVS-Net Target dominates molecule-only encoding even on chemical perturbations, indicating that target identity carries information about cell-level effect that molecular structure alone does not. The full encoder ablation, including representations across cell, genome, protein-sequence, and protein-structure modalities, is reported in Appendix Table 8.

**Table 1:** Pairwise regression loss (MSE) of inferred networks on a context-held-out split for each perturbation type. All rows are evaluated on the intersection of held-out perturbations for fair comparison. Per-modality choice of biochemical encoder upstream of the context encoder, and the full encoder ablation, are reported in Appendix C and Table 8.

| Model | Chemical | shRNA | Over-Expression | Ligand |
| --- | --- | --- | --- | --- |
| Population | 1.0594 | 0.9788 | 0.7769 | 0.9315 |
| CellVS-Net Molecule | 0.6003 | — | — | — |
| CellVS-Net Target | **0.5633** | **0.6740** | **0.6801** | **0.5873** |

### Training on Disjoint Modalities Improves Overall Performance

The four perturbation modalities in LINCS L1000 (chemical, shRNA, over-expression, ligand) act on shared cellular machinery but are only observed one at a time and are typically modeled in isolation. A single CellVS-Net context encoder accepts heterogeneous context features and supports joint training across all four, allowing dependency structure learned from one modality to inform predictions for the others. We train one joint encoder on the union of all perturbation types using the best-performing context representation for each modality from Table 1 and compare against the per-modality encoders (Table 2). We find that joint training across contexts, even for disjoint context modalities, reduces network MSE on every modality. We see the largest gains on the data-poorer genetic perturbations: shRNA improves from 0.674 to 0.552 (-18%) and over-expression from 0.680 to 0.341 (-50%). Chemical and ligand modalities, which already have larger training sets, see smaller but consistent improvements (-2% and -8%). This pattern is consistent with positive transfer from data-rich to data-scarce modalities through a shared dependency-network output space.

**Table 2:** Network prediction MSE using ChemBERTa [31] representations for chemical perturbations and gene representations for other perturbation types (from Table 1) compared against CellVS-Net trained on all perturbations together with those same representations.

| Model | Chem | shRNA | Over-Expression | Ligand |
| --- | --- | --- | --- | --- |
| CellVS-Net (separate) | 0.6003 | 0.6740 | 0.6801 | 0.5873 |
| CellVS-Net (joint) | 0.5907 | 0.5524 | 0.3410 | 0.5419 |

### Drug-Disease Retrieval: Predicting Disease Indications for Drugs with Novel Targets

A useful cell-level screen should cluster drugs with similar therapeutic effects even when they hit different targets. We assemble small-molecule drugs from OpenTargets with disjoint target profiles but a shared FDA-approved disease, providing a ground truth that target-centric methods cannot recover by construction. CellVS-Net Target does not apply in this target-disjoint setting. We apply CellVSNet Molecule and train baseline expression models in the same setting using identical chemical input representations. CellVS-Net Molecule networks place same-disease drugs closer than expression, foundation-model embeddings, target-binding, or fingerprint baselines (Table 3; Figure 4). Critically, CellVS-Net is the only representation whose Hits@5 is significantly different from random after multiple-testing correction (Table 4); it also surpasses the oracle expression upper bound, indicating that gene-network restructuring carries therapeutic signal that raw expression snapshots do not. The clustermap in Figure 4 visualizes this: same-disease drugs form more cohesive blocks under network distance, whereas expression-based distances mix across indications.

**Table 3:**
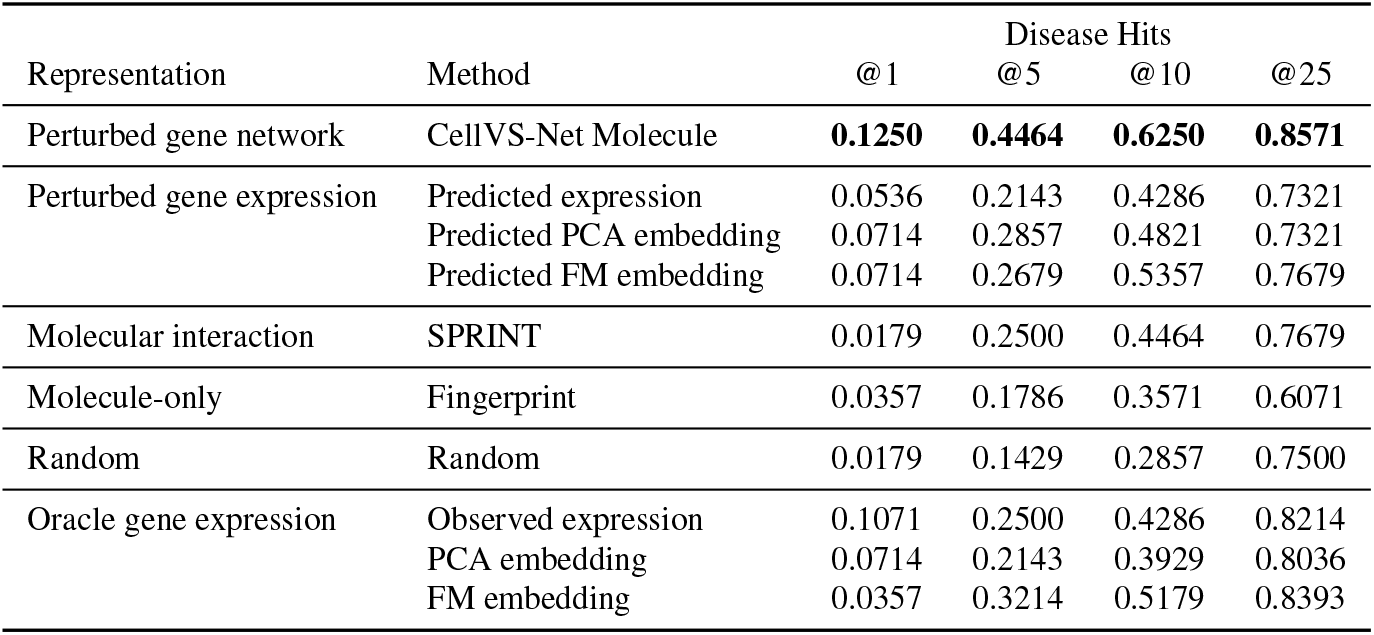
Evaluating methods of representing small molecule drugs in terms of their ability to predict FDA approvals. We compile a dataset of diseases, targets, and small molecules, where each disease has multiple approved small molecule drugs targeting different genes or sets of genes. We hold out drugs with identical target profiles, and use each held-out drug to query the remaining drugs, returning the *k* nearest neighbors in terms of Euclidean distance with *k* ∈ {1, 5, 10, 25}. We report a hit if any of the returned drugs have an FDA approval for the same disease as the held-out drug. Paired bootstrap p-values for Hits@5 versus the random baseline are shown in Table 4; 95% confidence intervals on every cell are listed in the appendix Table 11.

**Table 4:**
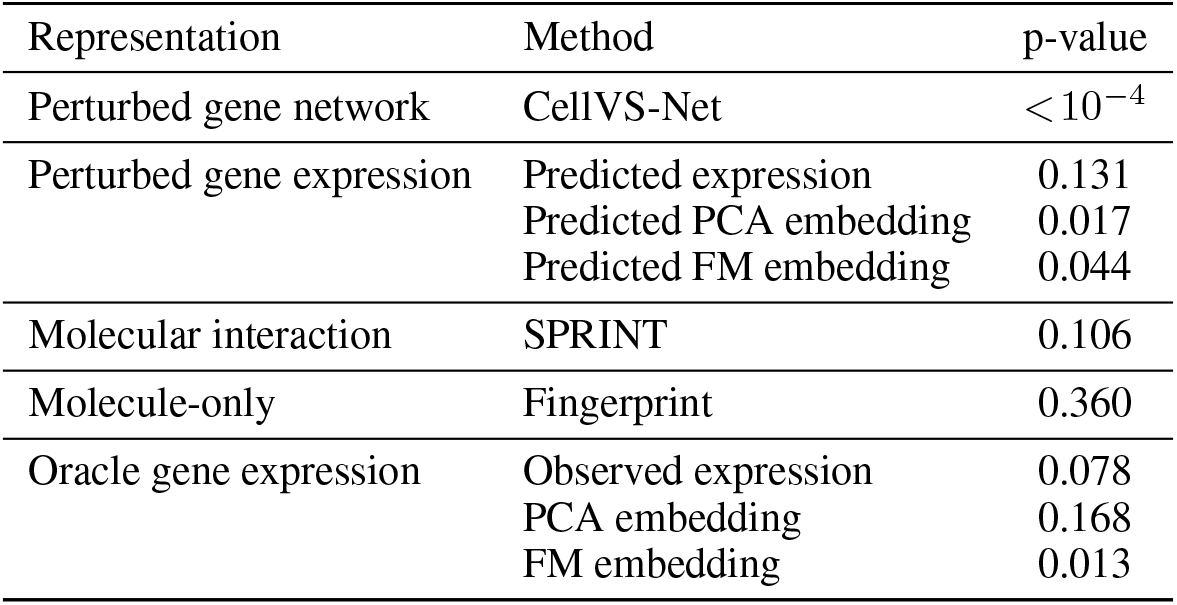
Paired bootstrap p-values (10,000 resamples) for Hits@5 versus random performance on DDR-Bench. Row ordering matches Table 3. CellVS-Net is the only method significantly better than random after multiple testing correction.

| Representation | Method | p-value |
| --- | --- | --- |
| Perturbed gene network | CellVS-Net | $< 10^{-4}$ |
| Perturbed gene expression | Predicted expression | 0.131 |
|  | Predicted PCA embedding | 0.017 |
|  | Predicted FM embedding | 0.044 |
| Molecular interaction | SPRINT | 0.106 |
| Molecule-only | Fingerprint | 0.360 |
| Oracle gene expression | Observed expression | 0.078 |
|  | PCA embedding | 0.168 |
|  | FM embedding | 0.013 |

**Figure 4:**
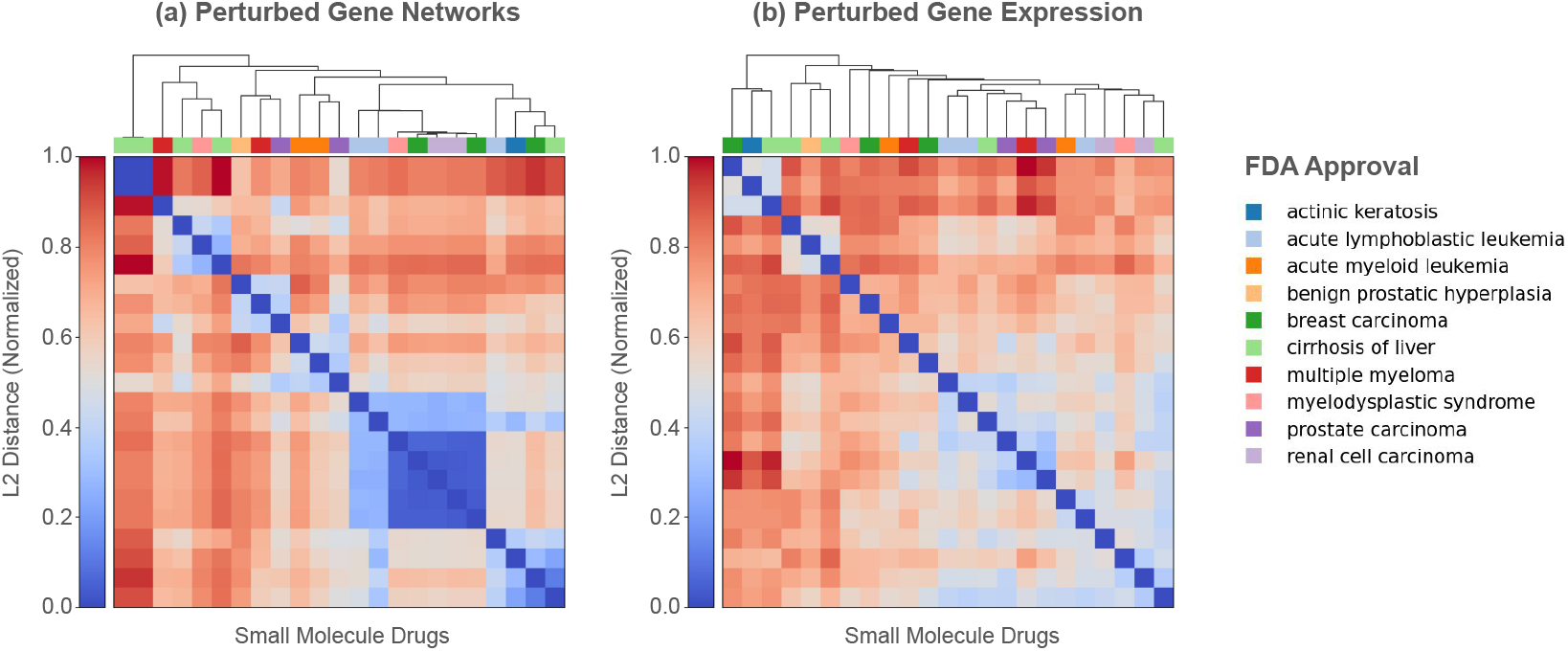
Organization of drugs based on (a) perturbed gene networks and (b) perturbed gene expression representations. Drugs are annotated with their FDA-approved disease indications. All samples are taken from the PC3 cell type.

### Drug-Target Retrieval: Matching Synonymous Perturbations Across Modalities

DTR-Bench asks whether known drug-target interactions can be reconstructed from cell-level signatures, bridging cell-level screens to traditional drug-target interaction prediction. We pair LINCS chemical perturbations with shRNA knockdowns of their known targets. We apply CellVS-Net Molecule to encode chemical perturbations and CellVS-Net Target to encode knockdowns. We train baseline expression models in the same setting using identical chemical and target input representations. We also compare to SPRINT [1], a SOTA target-binding model that does not see cellular readout. CellVS-Net outperforms all oracle and predicted baselines on global drug-target graph reconstruction, and is the only method whose AUROC and AUPRC are significantly above random (Table 5). SPRINT, which is directly optimized for drug-target binding from molecular structure, performs at chance on this protocol. This gap motivates representations that act on shared cellular machinery rather than on molecular structure alone. Per-query lookup results and a web tool for browsing shRNA - chemical embeddings are in Appendix Table 13 and Appendix Figure **??**.

**Table 5:** Recovering known drug-target relationships using different perturbation representations on DTR-Bench. AUROC and AUPRC are calculated using ground-truth and predicted bipartite drug-target graphs via distance thresholding. Expression-based representations are derived from LINCS L1000; PCA applies a 50-component PCA to the full dataset; AIDO.Cell embeds each sample. One-sided paired bootstrap p-values versus the random baseline are reported alongside each metric (10,000 resamples); 95% confidence intervals are listed in appendix Table 12.

|  | AUROC | p-value | AUPRC | p-value |
| --- | --- | --- | --- | --- |
| CellVS-Net predicted networks | **0.540** | **0.0013** | **0.012** | **0.0028** |
| Predicted expression | 0.482 | 0.639 | 0.009 | 0.430 |
| Predicted PCA embedding | 0.476 | 0.758 | 0.009 | 0.268 |
| Predicted FM embedding | 0.501 | 0.261 | 0.009 | 0.121 |
| SPRINT | 0.445 | 0.993 | 0.008 | 0.874 |
| Random | 0.489 | — | 0.008 | — |
| Oracle observed expression | 0.513 | 0.082 | 0.009 | 0.132 |
| Oracle PCA embedding | 0.521 | 0.033 | 0.010 | 0.065 |
| Oracle FM embedding | 0.521 | 0.027 | 0.010 | 0.022 |

## Discussion

Cell-level screening methods promise to move late-stage drug-discovery failures into earlier *in silico* screening stages, but have been insufficiently validated thus far. In this study, we provide the first quantitative benchmarks for cell-level virtual screening through the formulation of the DDR-Bench and DTR-Bench benchmarks. Both benchmarks are modality-agnostic, evaluating drug, drug-target, and cell-level modeling approaches across several data modalities on practical drug discovery tasks: recovering disease indications for drugs with unseen targets (Table 3) and reconstructing drug-target relationships from the effects of different perturbation modalities (Table 5). These benchmarks reveal the gaps in the performance of current molecular and transcriptomic methods.

To address this, we develop CellVS-Net, a highly programmable method for predicting how gene networks restructure under different cell types, small molecule therapeutics, signaling molecules, gene knockdowns, and gene over-expressions. CellVS-Net introduces a scalable differentiable surrogate loss for multivariate Gaussian likelihood that decomposes covariance estimation into pairwise regressions, with desirable statistical properties for estimation in long-tail drug screening applications (Table 14). The result generalizes smoothly to unseen drugs, doses, and cell types, reducing MSE by 12–47% across modalities (Table 1). The surrogate loss is a drop-in replacement for the isotropic-Gaussian (mean squared error) loss that dominates current cell-modeling pipelines, and adds context-specific gene-gene dependence as an output without changing the encoder architecture.

When applied zero-shot to DDR-Bench and DTR-Bench, CellVS-Net improves over molecular and expression baselines, including the oracle expression upper bound that represents an idealized expression prediction method (Tables 3, 4, 5). The fact that gene networks surpass this oracle indicates that structured representations of gene-gene dependence carry therapeutically relevant signal that is not recoverable from raw expression snapshots, even in principle. This finding redirects the virtual-cell research agenda away from reconstruction accuracy alone and toward downstream therapeutic utility as a primary evaluation criterion.

## Limitations

Overall, cell-level virtual screening has potential to shift drug-discovery effort from late-stage *in vitro* and *in vivo* assays toward earlier *in silico* prioritization, reducing experimental cost and animal use and supporting rejection of candidates with system-level efficacy or toxicity issues before any new assay is run. However, there are several limitations with the current work. CellVS-Net does not predict human-subject outcomes directly and is not a substitute for preclinical or clinical safety evaluation. The work here develops CellVS-Net as a proof of concept, not a production-ready tool. Baselines were chosen based on current SOTA, common practice, and alignment with the zero-shot representation evaluation setting. Current cell-level evaluations are limited to expression-based methods due to the availability of oracle data, but cell imaging should be considered in future work. The current version of CellVS-Net was trained on PCA-compressed expression data due to memory constraints. On downstream DDR-Bench retrieval, joint training of CellVS-Net degrades chemical-perturbation Hits@*k* relative to a chemical-only encoder (Appendix Table 10), suggesting that better network reconstruction does not strictly translate to better drug retrieval and that the per-modality encoder remains preferable when the screening task is itself modality-specific. On DTR-Bench, labels are inferred by matching all drugs with knockdown perturbations. This matching method assumes all drugs are inhibitors, the most common drug class, but will result in label noise for cell-level evaluations on non-inhibitory drugs. In the future, more sophisticated matches to knockdowns, over-expressions, and ligand perturbations can be made based on mechanism of action.

## A Resources

We provide source code for data curation, model training, and evaluations at https://github.com/SohanAddagudi/contextpert.

## B Training

For each perturbation type in the LINCS L1000 dataset (small molecule, shRNA, over-expression, ligand) we apply quality control filters based on replicate correlation and self-ranking performance to ensure high-confidence perturbation profiles, then hold-out 20% of perturbations at random. We construct a context vector *C* for each sample from metadata including perturbation type, target gene (for genetic perturbations), dose, timepoint, and control expression for the corresponding cell line. Expression measurements are compressed to 50 metagenes using principal component analysis, inferred from the train set. All contexts and expression samples are feature-normalized according to train-set mean and standard deviation prior to fitting. To train the model, we apply the Contextualized modeling Python library [32]. We test several methods for representing perturbations to improve generalization to unseen conditions, described in the section below.

### Compute resources

Each CellVS-Net training run reported in Tables 8 and 15 takes approximately 1h on a single NVIDIA H100 80GB GPU. Pre-computing perturbation embeddings runs once per representation: AIDO.Cell embeddings for all samples take roughly 24h, while each of the other gene-level embeddings used as context in Table 8 (AIDO.DNA, AIDO.Protein, AIDO.StructureTokenizer, ChemBERTa, PCA) takes roughly 1h, and SPRINT embeddings used in Tables 3 and 13 take roughly 1h. End-to-end data processing from raw LINCS L1000 and OpenTargets sources takes about 2h on CPU with 16GB memory and ∼200GB of disk space; all remaining benchmark scripts (table generation, retrieval, bootstrap intervals) complete in under 5min of CPU time.

## C Perturbation Representations

We employ multiple representation strategies for perturbations in our trained networks, motivated by recent efforts in benchmarking multimodal foundation models for cellular perturbation prediction [33]. For small-molecule perturbations, we use SMILES-based molecular representations, while for all perturbation types, we also explore target-based representations derived from gene-level embeddings.

### SMILES-based networks

For small-molecule perturbations, we compare two chemical featurization strategies. First, we compute Morgan fingerprints [24], a substructure representation that encodes local atomic environments and has proven effective in traditional cheminformatics pipelines. Second, we apply ChemBERTa-100M-MLM [31], a transformer-based molecular foundation model trained on large SMILES corpora, which provides contextualized embeddings that better capture semantic and structural relationships among compounds. These two representations provide complementary baselines for evaluating molecular embedding quality and their effect on drug–target inference.

### Target-based networks

For perturbations with gene targets, we integrate embeddings from multiple biological foundation models spanning expression, genomic sequence, and protein structure modalities.

### AIDO.Cell (expression-based)

We use AIDO.Cell 100M [34], a full-transcriptome single-cell foundation model trained across diverse cellular contexts. Gene embeddings are computed using K562 control cells from Norman et al [35].

### AIDO.DNA (sequence-based)

We extract sequence-level gene representations using the AIDO.DNA model [36]. For each gene, we define a 4 kbp window centered at the transcription start site (TSS), run model inference to obtain nucleotide embeddings, and apply mean pooling across the sequence to generate a single fixed-length embedding vector per gene.

### AIDO.Protein (structure-informed)

To capture protein-level information, we utilize AIDO.ProteinIF-16B [37], a large-scale model trained jointly on sequence and inferred structure representations. Residue-level embeddings are mean-pooled to yield protein-level embeddings, and for genes encoding multiple isoforms, we average across all available proteins.

### AIDO.StructureTokenizer (geometry-based)

We further incorporate 3D structural information using the AIDO.StructureTokenizer model [38], which tokenizes protein backbone geometry and side-chain orientations to produce structure-aware embeddings. For genes with multiple resolved structures, we mean-pool over all available embeddings.

### PCA (non-FM)

As described in previous benchmarking studies [33], we derive baseline gene embeddings by applying PCA to control-condition expression profiles. For each gene, we collect its unperturbed expression values across all control samples and project this vector into a PCA space learned over the full control expression matrix (compressing variation across samples). This was once again computed with K562 control cells from Norman.

## D Prediction of Molecular Representations

We trained supervised regression models to predict perturbation-induced molecular representations from chemical structure–derived embeddings.

### Input Features

For all prediction tasks, the input representation *X* ∈ ℝ^*d*^ consisted of precomputed chemBERTa embeddings associated with the compound corresponding to each perturbation instance. These embeddings are fixed-length continuous vectors derived from SMILES strings and are independent of cellular context.

### Predictor Model

We used a multi-output ridge regression model to map chemical embeddings to molecular representations. Given an input matrix *X* ∈ ℝ^*n*×*d*^ and target matrix *Y* ∈ ℝ^*n*×*p*^, the model solves

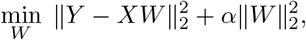

where *W* ∈ ℝ^*d*×*p*^ is the regression weight matrix and *α* = 1.0 is the regularization parameter. A separate model was trained for each type of molecular representation. Models were fit using only perturbation instances in the training split and then used to generate predictions for all instances.

### Predicting PCA Metagenes

Gene expression profiles were first standardized and projected into a low-dimensional space using principal component analysis (PCA). The top *K* = 50 principal components were retained and treated as metagene features.

### Predicting Gene Expression

In the expression prediction setting, the supervision target *Y* consisted of the landmark gene expression vector for each perturbation instance.

### Predicting AIDO Cell 3M Embeddings

For representation learning with foundation-model embeddings, the supervision target was the 128-dimensional AIDO Cell 3M embedding associated with each perturbation instance.

## E Benchmark Curation

### DDR-Bench construction

We filter the combined OpenTargets-LINCS dataset to small molecules represented in LINCS chemical perturbations, then restrict to diseases with at least 2 Phase-IV FDA-approved drugs whose target signatures (sorted lists of Ensembl protein IDs known to be bound) are distinct. For evaluation, one target signature is held out at a time and used to query the remaining drugs. Coverage by disease is reported in Table 7.

### DTR-Bench construction

We filter the combined OpenTargets-LINCS dataset to drug-target pairs in which the drug appears as a LINCS chemical perturbation and the target appears as a LINCS shRNA knockdown. We apply quality control filters to both modalities based on replicate correlation and self-ranking performance to retain only high-confidence perturbation profiles. Summary statistics are reported in Table 6.

**Table 6:**
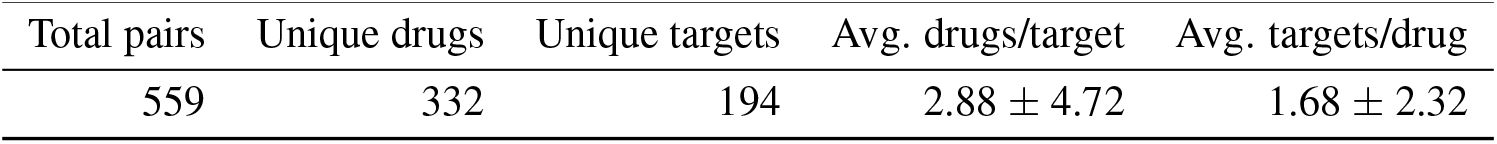
DTR-Bench summary statistics.

| Total pairs | Unique drugs | Unique targets | Avg. drugs/target | Avg. targets/drug |
| --- | --- | --- | --- | --- |
| 559 | 332 | 194 | $2.88 \pm 4.72$ | $1.68 \pm 2.32$ |

**Table 7:** DDR-Bench coverage by disease. For each disease, we report the number of distinct target signatures with at least one drug and the total number of distinct drugs mapped to those signatures. Target signatures are represented as a sorted list of Ensembl ids.

| Disease ID | Disease Name | Targets | Drugs |
| --- | --- | --- | --- |
| EFO_0000305 | breast carcinoma | 5 | 7 |
| EFO_0001422 | cirrhosis of liver | 5 | 5 |
| EFO_0000220 | acute lymphoblastic leukemia | 4 | 4 |
| EFO_0000222 | acute myeloid leukemia | 4 | 4 |
| EFO_0000284 | benign prostatic hyperplasia | 3 | 4 |
| EFO_0001378 | multiple myeloma | 3 | 4 |
| EFO_0002496 | actinic keratosis | 3 | 3 |
| EFO_0000681 | renal cell carcinoma | 3 | 3 |
| EFO_0000198 | myelodysplastic syndrome | 2 | 3 |
| EFO_0001663 | prostate carcinoma | 2 | 3 |
| EFO_1001469 | Mantle cell lymphoma | 2 | 2 |
| MONDO_0015760 | T-cell non-Hodgkin lymphoma | 2 | 2 |
| EFO_0004193 | basal cell carcinoma | 2 | 2 |
| EFO_1001012 | leptomeningeal metastasis | 2 | 2 |
| EFO_0004289 | lymphoid leukemia | 2 | 2 |
| EFO_1001051 | mycosis fungoides | 2 | 2 |
| EFO_0003060 | non-small cell lung carcinoma | 2 | 2 |
| EFO_1000045 | pancreatic neuroendocrine tumor | 2 | 2 |

## F Extended Results

### F.1 Encoder Ablation for CellVS-Net

**Table 8:** Pairwise regression loss (MSE) of inferred networks on a context-held-out split for various perturbation types. All CellVS-Net variants and the population baseline are evaluated on the intersection of all held-out perturbations for fair comparison. Best per column in bold.

| Model Type | Context Encoder | Chemical | shRNA | Over-Expression | Ligand |
| --- | --- | --- | --- | --- | --- |
| Population | None | 1.0594 | 0.9788 | 0.7769 | 0.9315 |
| CellVS-Net Molecule | Morgan Fingerprint [24] | 0.5786 | — | — | — |
|  | ChemBERTa [31] | 0.6003 | — | — | — |
| CellVS-Net Target | AIDO.Structure [38] | 0.5795 | 0.6741 | 0.6907 | **0.5873** |
|  | AIDO.Protein [37] | **0.5633** | **0.6740** | **0.6801** | 0.5890 |
|  | AIDO.Cell [11] | 0.6005 | 0.6754 | 0.7403 | 0.6414 |
|  | AIDO.DNA [36] | 0.6015 | 0.6817 | 0.7645 | 0.6808 |
|  | Gene PCA | 0.6105 | 0.6777 | 0.7070 | 0.6122 |

Context representations impose a prior on the similarity of downstream network estimation tasks for CellVS-Net. Good representations can greatly improve accuracy and generalization, even in the presence of noise features and non-linear effects in this modeling regime [25, 29]. We try several representations for small molecule therapeutics, signaling molecules, gene knockdowns, and gene over-expressions, aiming to produce a highly generalizable perturbation-specific network generator. We compare these context-adaptive models against a context-agnostic population estimator. Unlike previous experiments, group-specific modeling and one-hot contexts are not applicable in this regime, as unseen contexts cannot be mapped onto the original groups or feature set. We evaluate models in terms of the pairwise regression loss on held-out perturbations with expression measurements (Table 8).

CellVS-Net strongly outperforms the context-agnostic baseline by learning to map cell type and perturbation contexts to gene network rewiring. For small-molecule perturbations, CellVS-Net generalizes effectively to held-out molecules: representing molecules with structural fingerprints reduces error by 45%, and representing them by their known protein targets improves further to 47%. For shRNA, over-expression, and ligand perturbations, contexts are represented by their target gene or ligand protein. We include a non-pretrained context representation (Gene PCA) in each case to evaluate the importance of pretrained representations for generalization. Pretrained protein and structure representations consistently outperform PCA across genetic and ligand modalities, with AIDO.Protein achieving the lowest MSE on chemical, shRNA, and over-expression perturbations and AIDO.Structure on ligand perturbations.

### F.2 CellVS-Net Molecule Trained On All Available Drugs

**Table 9:**
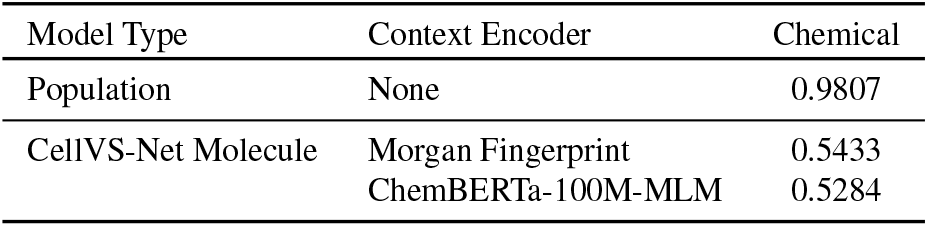
Mean squared error (MSE) of inferred networks across held out chemical perturbations. This evaluation uses the same test set as Table 1, but the training set includes all available drugs with corresponding SMILES strings, rather than only drugs with known targets.

### F.3 DDR-Bench Extended Results

**Table 10:** DDR-Bench drug-disease retrieval performance (from Table 3).

| Model | @ 1 | @ 5 | @ 10 | @ 25 |
| --- | --- | --- | --- | --- |
| CellVS-Net (separate) | 0.1250 | 0.4464 | 0.6250 | 0.8571 |
| Joint CellVS-Net (all types) | 0.0714 | 0.2857 | 0.5179 | 0.7857 |

**Table 11:** Evaluating methods of representing small molecule drugs in terms of their ability to predict FDA approvals. We compile a dataset of diseases, targets, and small molecules, where each disease has multiple approved small molecule drugs targeting different genes or sets of genes. We hold out drugs with identical target profiles, and use each held-out drug to query the remaining drugs, returning the *k* nearest neighbors in terms of Euclidean distance with *k* ∈ {1, 5, 10, 25}. We report a hit if any of the returned drugs have an FDA approval for the same disease as the held-out drug. Confidence intervals (95%) are computed via bootstrap (10,000 resamples) where available.

| Representation | Method | Disease Hits |  |  |  |
| --- | --- | --- | --- | --- | --- |
|  |  | @ 1 | @ 5 | @ 10 | @ 25 |
| Perturbed gene network | CellVS-Net | **0.1250 [0.0536, 0.2143]** | **0.4464 [0.3214, 0.5714]** | **0.6250 [0.5000, 0.7500]** | **0.8571 [0.7679, 0.9464]** |
| Perturbed gene expression | Predicted expression | 0.0536 [0.00, 0.13] | 0.2143 [0.11, 0.32] | 0.4286 [0.31, 0.55] | 0.7321 [0.61, 0.84] |
|  | Predicted PC | 0.0714 [0.02, 0.14] | 0.2857 [0.18, 0.41] | 0.4821 [0.36, 0.61] | 0.7321 [0.61, 0.84] |
|  | Predicted AIDO.Cell | 0.0714 [0.02, 0.14] | 0.2679 [0.16, 0.39] | 0.5357 [0.41, 0.66] | 0.7679 [0.64, 0.88] |
| Molecular interaction | SPRINT | 0.0179 [0.000, 0.054] | 0.2500 [0.143, 0.375] | 0.4464 [0.321, 0.571] | 0.7679 [0.661, 0.875] |
| Molecule-only | Fingerprint | 0.0357 [0.000, 0.089] | 0.1786 [0.089, 0.286] | 0.3571 [0.232, 0.482] | 0.6071 [0.482, 0.732] |
| Random | Random | 0.0179 [0.0000, 0.0536] | 0.1429 [0.0536, 0.2321] | 0.2857 [0.1786, 0.4107] | 0.7500 [0.6250, 0.8571] |
| Oracle gene expression | Observed expression | 0.1071 [0.0357, 0.1964] | 0.2500 [0.1429, 0.3571] | 0.4286 [0.3036, 0.5536] | 0.8214 [0.7143, 0.9107] |
|  | PCA expression | 0.0714 [0.018, 0.143] | 0.2143 [0.107, 0.321] | 0.3929 [0.268, 0.518] | 0.8036 [0.696, 0.911] |
|  | AIDO.Cell embedding | 0.0357 [0.000, 0.089] | 0.3214 [0.196, 0.446] | 0.5179 [0.393, 0.643] | 0.8393 [0.732, 0.929] |

### F.4 DTR-Bench Extended Results

**Table 12:** Recovering known drug–target relationships using different perturbation representations. AUROC and AUPRC are calculated using ground-truth and predicted bipartite drug–target graphs, using distance thresholding to induce predictions. Performance is evaluated on DTR-Bench with bootstrap confidence intervals (10,000 resamples; 332×194 pairs).

|  | AUROC (95% CI) | AUPRC (95% CI) |
| --- | --- | --- |
| CellVS-Net predicted networks | **0.5399 [0.5165, 0.5631]** | **0.0123 [0.0093, 0.0176]** |
| Predicted expression | 0.4822 [0.4572, 0.5070] | 0.0085 [0.0075, 0.0096] |
| Predicted PCA expression | 0.4763 [0.4509, 0.5015] | 0.0089 [0.0078, 0.0101] |
| Predicted AIDO.Cell embeddings | 0.5005 [0.4752, 0.5256] | 0.0093 [0.0082, 0.0106] |
| SPRINT [1] | 0.4447 [0.4206, 0.4691] | 0.0076 [0.0068, 0.0086] |
| Random | 0.4886 [0.4637, 0.5126] | 0.0084 [0.0075, 0.0094] |
| Oracle AIDO.Cell [11] | 0.5208 [0.4977, 0.5445] | 0.0104 [0.0087, 0.0130] |
| Oracle PCA expression | 0.5207 [0.4967, 0.5453] | 0.0096 [0.0084, 0.0111] |
| Oracle observed expression | 0.5130 [0.4890, 0.5377] | 0.0092 [0.0082, 0.0105] |

**Table 13:** Recovering known drug–target relationships using different perturbation representations. AUROC and AUPRC are calculated using ground-truth and predicted bipartite drug–target graphs, using distance thresholding to induce predictions. Query-level recall rates (Hits@*k*) are reported for both drug→target and target→drug retrieval tasks as Drug Hits and Target Hits respectively. Expression-based representations were derived from LINCS L1000 small molecule and shRNA data. PCA applies a 50-component PCA to this full dataset. AIDO.Cell embeds each sample.

| Method | AUROC | AUPRC | Drug→Target Hits |  |  |  | Target→Drug Hits |  |  |  |
| --- | --- | --- | --- | --- | --- | --- | --- | --- | --- | --- |
|  |  |  | @1 | @5 | @10 | @50 | @1 | @5 | @10 | @50 |
| CellVS-Net predicted networks | **0.5399**<br>(0.5165-0.5631) | **0.0123**<br>(0.0093-0.0176) | 0.0000<br>(0.0000-0.0000) | 0.0272<br>(0.0121-0.0453) | 0.0332<br>(0.0151-0.0544) | 0.3230<br>(0.2719-0.3716) | 0.0261<br>(0.0052-0.0518) | 0.0468<br>(0.0207-0.0777) | 0.0675<br>(0.0363-0.1036) | 0.3108<br>(0.2435-0.3782) |
| Predicted expression | 0.4822<br>(0.4756-0.5251) | 0.0085<br>(0.0079-0.0101) | 0.0150<br>(0.0030-0.0242) | 0.0480<br>(0.0121-0.0453) | 0.0781<br>(0.0453-0.1027) | 0.3463<br>(0.2779-0.3807) | 0.0155<br>(0.0000-0.0361) | 0.0464<br>(0.0206-0.0773) | 0.0875<br>(0.0206-0.0825) | 0.2269<br>(0.1701-0.2887) |
| Predicted PCA expression | 0.4763<br>(0.4633-0.5120) | 0.0089<br>(0.0078-0.0102) | 0.0179<br>(0.0000-0.0211) | 0.0510<br>(0.0302-0.0755) | 0.0781<br>(0.0514-0.1088) | 0.3281<br>(0.2628-0.3656) | 0.0154<br>(0.0000-0.0361) | 0.0362<br>(0.0103-0.0567) | 0.0670<br>(0.0412-0.1186) | 0.2010<br>(0.1959-0.3196) |
| Predicted AIDO.Cell embeddings | 0.5005<br>(0.4861-0.5344) | 0.0093<br>(0.0081-0.0108) | 0.0061<br>(0.0000-0.0211) | 0.0393<br>(0.0121-0.0514) | 0.0756<br>(0.0393-0.0906) | 0.2567<br>(0.2236-0.3172) | 0.0261<br>(0.0000-0.0258) | 0.0777<br>(0.0361-0.1031) | 0.1395<br>(0.0773-0.1649) | 0.3351<br>(0.2371-0.3660) |
| SPRINT [1] | 0.4447<br>(0.4206-0.4691) | 0.0076<br>(0.0068-0.0086) | **0.0332**<br>(0.0151-0.0544) | **0.1025**<br>(0.0695-0.1360) | **0.1507**<br>(0.1118-0.1903) | 0.3048<br>(0.2568-0.3565) | 0.0051<br>(0.0000-0.0155) | 0.0514<br>(0.0206-0.0876) | 0.0773<br>(0.0412-0.1186) | 0.2780<br>(0.2165-0.3402) |
| Random | 0.4886<br>(0.4637-0.5126) | 0.0084<br>(0.0075-0.0094) | 0.0030<br>(0.0000-0.0090) | 0.0331<br>(0.0151-0.0542) | 0.0630<br>(0.0392-0.0904) | 0.3253<br>(0.2771-0.3765) | 0.0052<br>(0.0000-0.0155) | 0.0155<br>(0.0000-0.0361) | 0.0414<br>(0.0155-0.0722) | 0.2734<br>(0.2113-0.3351) |
| Oracle AIDO.Cell [11] | 0.5208<br>(0.4977-0.5445) | 0.0104<br>(0.0087-0.0130) | 0.0151<br>(0.0030-0.0301) | 0.0484<br>(0.0271-0.0723) | 0.0634<br>(0.0392-0.0904) | 0.3135<br>(0.2651-0.3645) | 0.0206<br>(0.0052-0.0412) | 0.0927<br>(0.0515-0.1340) | 0.1236<br>(0.0773-0.1701) | 0.3451<br>(0.2784-0.4124) |
| Oracle PCA expression | 0.5207<br>(0.4967-0.5453) | 0.0096<br>(0.0084-0.0111) | 0.0120<br>(0.0030-0.0242) | 0.0422<br>(0.0211-0.0665) | 0.0723<br>(0.0453-0.1027) | **0.3384**<br>(0.2870-0.3897) | **0.0516**<br>(0.0206-0.0876) | **0.1136**<br>(0.0722-0.1598) | **0.1392**<br>(0.0928-0.1907) | **0.3718**<br>(0.3041-0.4383) |
| Oracle observed expression | 0.5130<br>(0.4890-0.5377) | 0.0092<br>(0.0082-0.0105) | 0.0151<br>(0.0030-0.0302) | 0.0332<br>(0.0151-0.0544) | 0.0757<br>(0.0483-0.1057) | 0.3293<br>(0.2779-0.3807) | 0.0207<br>(0.0052-0.0412) | 0.0569<br>(0.0258-0.0928) | 0.1084<br>(0.0670-0.1546) | 0.2943<br>(0.2320-0.3608) |

### F.5 The Effect of Context Encoding

#### F.5.1 Modeling in Low-Data Settings

Here, contexts are one-hot encoded, containing no prior knowledge of cell line similarity, intentionally disadvantaging contextualized networks, which must learn to share information and extrapolate between modeling tasks from scratch. We also strip away perturbations and focus only on control measurements for each cell line to isolate the role of context sharing.

**Table 14:** Mean-squared error (MSE) of inferred transcriptional networks on a sample-held-out split for control measurements from all cell lines. CellVS-Net and group-specific models use one-hot encoded celltype contexts. Full Test contains all held-out samples. *n*_*c*_ *>* 3 assesses conditions with more than 3 observations, while *n*_*c*_ ≤ 3 assesses conditions with less than 3 observations.

| | Full Test | $n_c > 3$ | $n_c \leq 3$ |
| --- | --- | --- | --- |
| Population | 0.9859 | 0.9860 | 0.9184 |
| Group-specific | 0.6680 | 0.6668 | 3.9306 |
| CellVS-Net | 0.6169 | 0.6169 | **0.7011** |
| + dose, time | **0.6065** | **0.6064** | 0.8912 |

Table 14 shows that CellVS-Net achieves the best performance on the full dataset by mitigating the failure modes of the population and condition-specific baselines. Population models suffer from high bias, underfitting due to their inability to model cell line-specific effects, while cell line-specific models dramatically overfit on conditions with few samples (*n*_*c*_ ≤ 3), with MSE exploding in low-sample regimes. In contrast, CellVS-Net automatically interpolates between a population-like default when data are scarce and cell line-specific behavior when sufficient data are available, yielding stable performance across data regimes that more closely resemble the long-tail distribution of a virtual screening atlas.

#### F.5.2 Continuous Contexts Improve Generalization

**Table 15:** MSE of inferred networks on a sample-held-out split for perturbed expression measurements. Perturbation contexts are one-hot encoded, while different encoding schemes are used for cell line contexts.

| Model Variant | Mean Squared Error |
| --- | --- |
| Population | 0.9735 |
| Group-specific | 4.795e04 |
| CellVS-Net onehot | 0.6304 |
| CellVS-Net + dose, time, celltype | **0.6122** |

Next, we evaluate the impact of richer context features that are essential for extrapolating to unseen conditions. Continuous covariates such as dose and time, or high-dimensional summaries of cell state, are difficult to incorporate into discrete group-based models, which typically require hand-crafted bins or separate models per group. In a virtual screening setting, however, new compounds will often be proposed at doses and timepoints that do not exactly match those in the training data, and any useful model must interpolate smoothly across these axes.

To study this, we move from control-only networks to prediction of post-perturbation networks and incrementally augment the input features of the context encoder (Table 15). We represent small-molecule identities with one-hot encodings and vary the representation of the cell-type context from a one-hot label to embeddings of the unperturbed transcriptomic profile. Post-perturbation prediction is more challenging than the control-only setup in Table 14, yet CellVS-Net again avoids extreme over- and under-fitting. Replacing one-hot cell-type indicators with control expression and augmenting with dose and time substantially improves generalization for predicting post-perturbation networks. These results support the view that rich, continuous context encodings are necessary for CellVS-Net to achieve the smooth extrapolation across doses, timepoints, and cell types that virtual screening requires.

### F.6 Organization of Drug Effects Across Methods and Cell Lines

**Figure 5:**
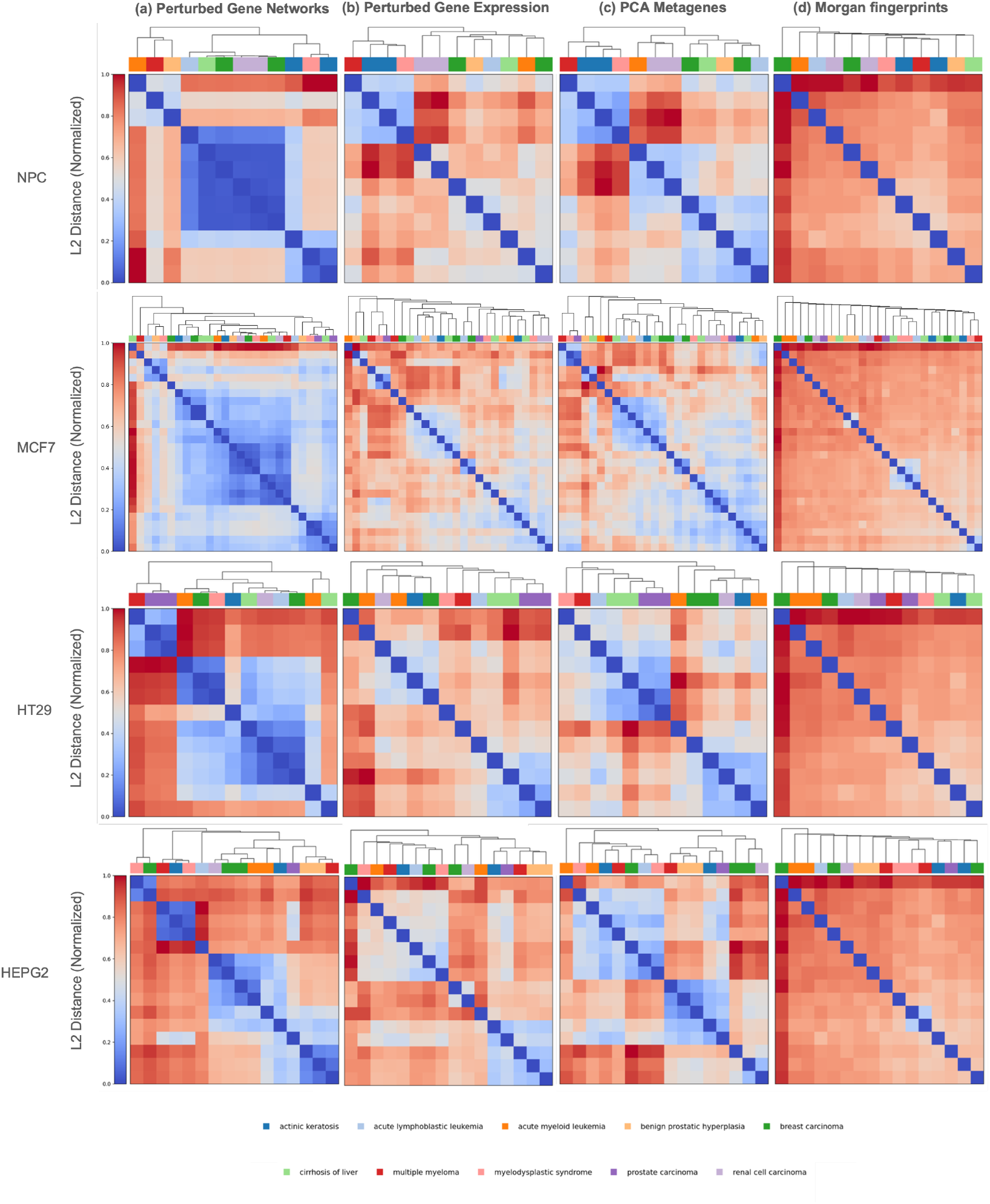
Organization of drugs based on four representations across cell types.

**Figure 6:**
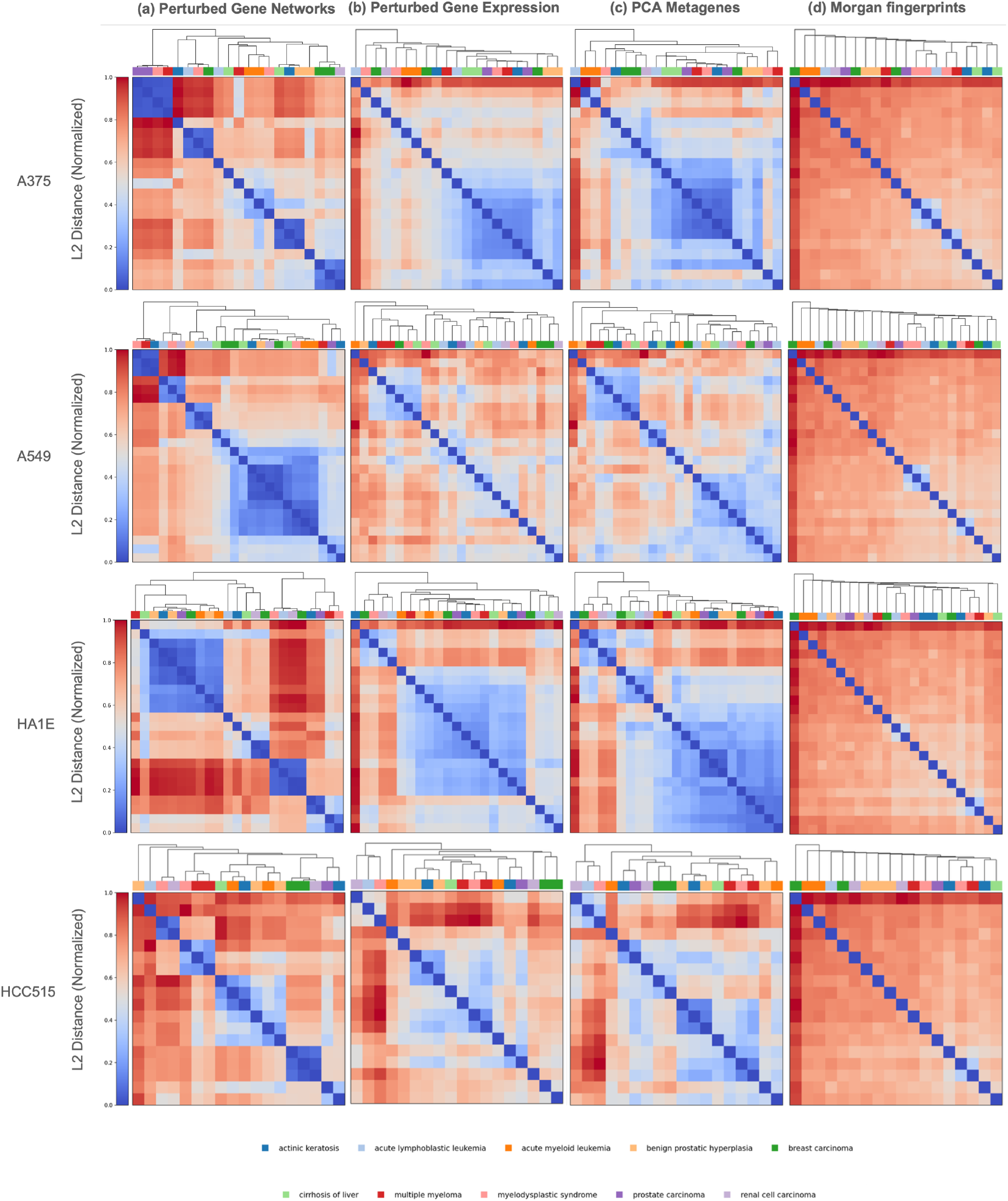
Organization of drugs based on four representations across cell types.

